# Soil microbial communities associated with giant sequoia: How does the world’s largest tree affect some of the world’s smallest organisms?

**DOI:** 10.1101/807040

**Authors:** Chelsea J. Carey, Sydney I. Glassman, Thomas D. Bruns, Emma L. Aronson, Stephen C. Hart

## Abstract

Giant sequoia (*Sequoiadendron giganteum*) is an iconic conifer that lives in relic populations on the western slopes of the California Sierra Nevada. In these settings it is unusual among the dominant trees in that it associates with arbuscular mycorrhizal fungi rather than ectomycorrhizal fungi. However, it is unclear whether differences in microbial associations extends more broadly to non-mycorrhizal components of the soil microbial community. To address this question we characterized microbiomes associated with giant sequoia and co-occurring sugar pine (*Pinus lambertiana*) by sequencing 16S and ITS1 of the bulk soil community at two groves with distinct parent material. We found tree-associated differences were apparent despite a strong grove effect. Bacterial/archaeal richness was greater beneath giant sequoia than sugar pine, with a unique core community that was double the size. The tree species also harbored compositionally distinct fungal communities. This pattern depended on grove but was associated with a consistently elevated relative abundance of *Hygrocybe* species beneath giant sequoia. Compositional differences between host trees correlated with soil pH, calcium availability, and soil moisture. We conclude that the effects of giant sequoia extend beyond mycorrhizal mutualists to include the broader community, and that some but not all host tree differences are grove-dependent.

## Introduction

There is increasing evidence that tree species influence a combination of soil chemical, physical, and biological properties (Hobbie *et al.*, 2007; Mitchell *et al.*, 2010; Langenbruch *et al.*, 2012). For example, variation in litter chemistry, patterns of nutrient uptake, root exudation, and microclimate among tree species can alter rates of decomposition, soil nitrogen (N) and carbon (C) availability, and pH (Binkley & Giardina, 1998). Tree-induced differences in resource availability and microclimate can, in turn, modify soil bacterial, archaeal, and fungal community composition (Ushio *et al.*, 2008; Prescott & Grayston, 2013) by encouraging microorganisms with compatible resource acquisition strategies and growth optima (Ayres *et al.*, 2009). In addition, under some circumstances, direct association with host-specific symbiotic microorganisms such as mycorrhizal fungi can further promote distinct soil fungal communities (Gao *et al.*, 2013; Urbanova *et al.*, 2015). Such plant-induced changes to microbial communities can become more or less pronounced with time since plant establishment (Strayer *et al.*, 2006), and can feedback on soil chemical and physical properties (Falkowski *et al.*, 2008), plant performance (Bever *et al.*, 2010; Aponte *et al.*, 2013), plant phenology (Wagner *et al.*, 2014), and plant community composition (van der Heijden *et al.*, 2008).

Although soil microorganisms can directly and indirectly influence plant dynamics (Abbott *et al.*, 2015) and may mediate how plant communities respond to anthropogenic threats (Malcolm *et al.*, 2006; Stephens *et al.*, 2014), information on soil microbial communities associated with many rare or endemic tree species is limited. One such tree is the giant sequoia (*Sequoiadendron giganteum*)—a species that epitomizes charismatic mega-flora (Hall *et al.*, 2011). Endemic to the western slope of the Sierra Nevada (California, USA), giant sequoia harbor a number of traits that make it unique, ecologically interesting, and of conservation concern. Most conspicuously, giant sequoia are the world’s largest trees, growing up to 87 m in height and reaching a bole volume of 1,500 m^3^ (Stephenson, 2000). They are also one of the longest-lived trees, with an estimated average age of 2,230 years and a maximum known age of 3,266 years (Stephenson, 2000). Over the last century, fire suppression has threatened giant sequoia regeneration by minimizing canopy gaps and exposure of mineral soil, two important factors for germination of this shade intolerant tree (York *et al.*, 2011). Future conditions marked by increased and prolonged drought are expected to put additional stress on giant sequoia (Su et al. 2017). Fortunately, their charisma has deemed them one of the seven natural wonders of the United States (DeFries, 2013), and together the ecological and cultural importance of the giant sequoia has promoted their current protection by both state and national agencies (Leisz, 1994; Lindenmayer *et al.*, 2014).

Despite their iconic status, very little is known about how giant sequoia influence the soil and even less is known about how these titans of the tree world interact with some of the smallest yet most important organisms on the planet: microorganisms. The few prior studies that have been conducted suggest that, similar to other members of the *Cupressaceae* family (Cross & Perakis, 2011a,b), giant sequoia accumulate relatively large amounts of base cations such as calcium in their leaf litter resulting in high soil base saturation compared to other mixed-conifer species (Zinke, 1962; Zinke & Stangenberger, 1994). In those same studies, giant sequoia was also reported to maintain relatively high values of soil pH and soil organic matter. From a microbial perspective, giant sequoia are known to associate with arbuscular mycorrhizal fungi (AMF), symbiotic fungi that form relationships with the vast majority of herbaceous plants (Fahey *et al.*, 2012). This contrasts with most other trees in the Sierra Nevada, such as those of the Pinaceae and the Fagaceae families, whose woody roots instead associate with symbiotic ectomycorrhizal fungi (EMF; Brundrett & Tedersoo, 2018). Few studies have probed the mycorrhizal dynamics of giant sequoia (Kough *et al.*, 1985; Molina, 1994; Fahey *et al.*, 2012), and none have intensively assessed soil bacterial/archaeal and fungal community structure (composition and diversity) using molecular techniques.

Given that giant sequoia occur on soils derived from various rock types—including granite, diorite, and andesite (Weatherspoon, 1990)—it is important to evaluate whether microbial dynamics beneath giant sequoia remain consistent or if they vary across groves with different parent material. Parent material exerts a strong influence on soil properties, and differences in underlying geology often interact with trees to shape soil microbial community structure (Ulrich & Becker, 2006; Carletti *et al.*, 2009; Wagai *et al.*, 2011). For example, parent material and vegetation type interacted to affect soil macroaggregate size, and both factors also shaped microbial community structure following 30 years of surface exposure of reclaimed surface mining sites (Yarwood *et al.*, 2015). The degree to which this occurs with the iconic giant sequoia, however, remains unknown.

In this paper, we sequenced bacterial, archaeal, and fungal communities from beneath giant sequoia and a co-dominant ectomycorrhizal tree, sugar pine (*Pinus lambertiana* Dougl.). Sugar pine are prevalent on the western slope of the Sierra Nevada and are second only to giant sequoia in total volume, with individuals reaching 76 m in height and living up to 600 years (Hardin *et al.*, 2001). We sampled soil from 32 individuals of each tree species across two groves with contrasting geologic substrates in Yosemite National Park, USA. By comparing these two tree species within and between groves, our experimental design allows us to evaluate for the first time the relative impact of these tree hosts and parent material on soil microbial structure. Using subsamples of soil collected for physiochemical analysis in a companion study (Hart et al., unpublished data), we sought to address the following questions: (Q1) what bacterial, archaeal, and fungal members comprise the soil microbial community beneath giant sequoia individuals?; (Q2) how do these microbial communities compare to those beneath co-occurring sugar pine individuals?; (Q3) are giant sequoia and sugar pine-associated microbial communities consistent across groves with differing geological substrates?; and (Q4) which soil characteristics, if any, correlate with tree-associated changes in microbial richness and composition?

## Methods

### Site Description

We sampled from two of the three giant sequoia groves contained within Yosemite National Park (YNP): Mariposa Grove (N 37.51539°, W 119.60435°) and Merced Grove (N 37.74982°, W 119.84061°). Both groves are located on the western slope of the Sierra Nevada in the rain-snow transition zone, and experience a Mediterranean-type climate with warm-dry summers and cool-wet winters. In addition to giant sequoia and sugar pine, other common trees in these groves are the ectomycorrhizal (EM) trees white fir (*Abies concolor*) and ponderosa pine (*Pinus ponderosa*), and the arbuscular mycorrhizal (AM) tree incense cedar (*Calocedrus decurrens*; Wang & Qiu, 2006, Allen & Kitajima, 2014). Both groves have relatively sparse understory vegetation composed primarily of AM-associated *Ceanothus* spp. and broadleaf lupine (*Lupinus latifolius*), as well seedlings and saplings of the EM-associated white fir (*Abies concolor*). Soils in the Mariposa Grove are derived from residuum and colluvium of metavolcanic (andesite and hornfels) with minor amounts of intermediate granitoid rock. Soils in the Merced Grove are derived from residuum and colluvium of quart-rich metasedimentary rock (USDA NRCS 2007). Despite contrasting parent materials, soils from both sites are classified within the same Soil Taxonomic Family: coarse-loamy, isotic, frigid Ultic Haploxeralfs.

### Experimental Design

Within each grove, we sampled from beneath 16 mature giant sequoia individuals and 16 co-occurring mature sugar pine individuals. Because mature giant sequoia trees were generally the limiting experimental unit (especially in the Merced Grove), we first selected mature giant sequoia individuals whose crowns did not overlap with adjacent trees. We then selected the closest mature sugar pine trees to each giant sequoia individual that shared similar aspect, slope, landscape position, and understory species (when present). Relatively little understory vegetation occurred beneath each focal tree, and areas containing nitrogen-fixing species such as *Ceanothus* spp. were avoided. As with giant sequoia, we ensured selection of sugar pine individuals whose crowns did not overlap with adjacent trees. This selection procedure was designed to minimize any confounding influences that may affect soil properties besides tree species. Our experimental design resulted in 32 total giant sequoia-sugar pine pairs that were typically within 30 m of each other.

### Soil Sampling

In August 2013, we sampled bulk surface soil from beneath each tree individual (i.e., we did not directly target rhizosphere soil surrounding plant roots). We selected sampling locations mid-crown and downslope of the tree bole with the assumption that aboveground litter would accumulate most at these locations and therefore the influence of trees would be maximal (Zinke, 1962). Five replicate soil cores (0-5 cm depth of mineral soil) per tree were taken within an approximately 20 cm x 20 cm area using an Oakfield corer (1.9 cm diameter; Oakfield Apparatus Co, Fond du Lac, WI, USA), and composited into a single sample within a sterile plastic bag (Whirl-Pak®, Nasco, Fort Atkinson, WI, USA). The soil corer was sanitized after each composite sample using a rinse of 10% bleach followed by a rinse of 95% ethanol. Soil samples were stored at 4°C, transported to University of California (UC) Merced, sieved (< 2 mm; sanitized between samples as described above), and subsampled for microbial analysis.

### DNA extraction

Each subsample was extracted immediately upon returning to the laboratory (within 24 hours), and two separate DNA aliquots per subsample were stored at −20 °C for subsequent analysis. Specifically, we extracted microbial DNA from 0.250 g soil (± 0.025) using a MO BIO PowerSoil Isolation Kit (Mo Bio Laboratories, Inc., Carlsbad, CA, USA) following the manufacturer’s instructions. One 50 µ L DNA aliquot was shipped on dry ice to UC Riverside for bacterial/archaeal 16S analysis and one was delivered on ice to UC Berkeley for fungal ITS analysis.

At the same time that DNA was extracted, additional soil subsamples were taken for 22 physiochemical analyses (Hart et al., unpublished data). These analyses were conducted on fresh or air-dried soils, as appropriate for the assay (Supplemental Information Methods S1).

### 16S rRNA amplicon preparation

After quantifying the extracted DNA using a NanoDrop 2000 (Thermo Fisher Scientific Inc., Wilmington, DE, USA), we amplified each sample in duplicate using primers targeting the V3-V4 region of the 16S rRNA gene (S-D-Bact-0341-b-S-17 and S-D-Bact-0785-a-A-21; Klindworth *et al.*, 2012). We conducted polymerase chain reaction (PCR) by combining 2.5 µL DNA template, 5 µL each of 1 µM forward and reverse primers, and 12.5 µL KAPA HiFi HotStart ReadyMix (KAPA Biosystems, Inc., Wilmington, MA, USA), totaling a 25 µL reaction. Thermocycler conditions were: 95 °C for 3 min., followed by 25 cycles of 95 °C for 30 s, 55 °C for 30 s, 72 °C for 30 s, followed by an extension step for 5 min. at 72 °C. After amplification, we combined and purified the duplicate PCR products using Agencourt AMPure XP Beads (Beckman Coulter Genomics, Danvers, MA, USA). A second round of PCR was subsequently conducted to attach dual indices to each sample using the Nextera XT Index Kit (Illumina Inc., San Diego, CA, USA). Briefly, 5 µL DNA, 5 µL each of 1 µM forward and reverse index primers, 25 µL KAPA HiFi HotStart ReadyMix, and 10 µL PCR grade water were combined to create a 50 µL mixture. Thermocycler conditions were: 95 °C for 3 min., followed by 8 cycles of 95 °C for 30 s, 55 °C for 30 s, 72 °C for 30 s, followed by an extension step for 5 min. at 72 °C. We then conducted a second AMPure purification step on the indexed amplicons, and quantified the products using the Quant-iT^TM^ PicoGreen^®^ dsDNA assay kit (Life Technologies Inc., Grand Island, NY, USA). As a final step, we pooled the samples together in equimolar concentrations and sequenced them in one Illumina MiSeq PE 2×300 run at the UC Riverside Genomics Core Facility, Riverside, CA, USA.

### ITS1 amplicon preparation

We PCR amplified the ITS1 spacer, a part of the internal transcribed spacer region (ITS) that serves as the universal DNA barcode for fungi, from each sample using the ITS1F-2 primer pair with Illumina MiSeq primers designed by Smith and Peay (2014). We conducted PCR by combining 1 µl DNA template, 0.50 µl of 10 µM forward primer, 1 µl of the 10µM bar-coded reverse primer, 0.130 µl of HotStar Taq Plus (5 units/µl) DNA polymerase (Qiagen, Valencia, CA, USA), 2.5 µl of 10 x PCR buffer supplied by the manufacturer, and 0.50 µl 10mM each dNTPs; the 25 µl reaction was brought to volume with water. Thermocycler conditions were: 95 °C for 5 min., followed by 29 amplification cycles of 94 °C for 30 s, 51 °C for 30 s, 72 °C for 1 min., followed by a 10 min. final extension at 72 °C. Amplifications for each barcoded sample were cleaned with Agencourt AMPure XP Beads, quantified with Qubit, pooled to equimolar concentration, and quality checked as described previously (Glassman *et al.*, 2016). The library was sequenced with Illumina MiSeq PE 2×250 at the Vincent J Coates Genomic Sequencing Laboratory, UC Berkeley, Berkeley, CA, USA.

### 16S Sequence analysis

We obtained the 16S rRNA gene sequences already demultiplexed from the UC Riverside Genomics Core Facility and processed them using Quantitative Insights into Microbial Ecology (QIIME; Caporaso *et al.*, 2010). After we joined the forward and reverse reads (allowing for 20% maximum difference within the region of overlap), we used default parameters to conduct quality control: reads were excluded if the length was less than 75 bases, if there were more than three consecutive low-quality base calls, if less than 75% of the read length was consecutive high-quality base calls, if a Phred score was below three, or if one or more ambiguous calls were present (Bokulich *et al.*, 2013). After quality filtering, 3.9 M sequence reads remained.

Operational taxonomic units (OTU) with 97% similarity were picked using open reference UCLUST against the 13_8 release of the Greengenes database (DeSantis *et al.*, 2006). Reads that did not match any sequences in the database were clustered *de novo* and singletons were filtered out. The median number of sequences per samples was 60,494, the mean was 61,422, and the range was from 18,378 to 114,657 sequences per sample. After removing unassigned OTUs (1% of sequences), the OTU table was rarefied to 36,345 reads per sample, and as a result, one sugar pine sample was dropped.

### ITS Sequence analysis

We obtained the ITS gene sequences already demultiplexed from the UC Berkeley Vincent J Coates Genomic Sequencing Laboratory and processed them as in (Glassman *et al.*, 2016) using UPARSE (Edgar, 2013). We removed distal priming/adapter sites, trimmed the remaining untrimmed, low-quality regions from the ends, and then joined the forward and reverse reads. Paired reads were then quality filtered using the fastq_filter command in usearch and employing a maximum expected number of errors of 0.25. After quality filtering, 2.9 M sequence read pairs remained. We picked 97% OTUs, then reference based chimera detection was employed using usearch and referencing against the UNITE database accessed on 10.09.2014 (Kõljalg *et al.*, 2005). After making the OTU table in usearch, we assigned taxonomy in QIIME (Caporaso *et al.*, 2010) using the same UNITE database. The resulting OTU table yielded 2,173 total OTUs. The median number of sequences per samples was 41,801 the mean was 39,946, and the range was from 4 to 75,202 sequences per sample. After removing all OTUs with a No Blast Hit, the OTU table was rarefied to 13,904 reads per sample; as a result, one giant sequoia and two sugar pine samples were dropped.

### Statistical Analysis

We used a multifaceted approach to assess microbial community structure of giant sequoia soils and to compare the structure of these communities with those beneath sugar pine. We first tested the main and interactive effects of plant species and grove on bacterial/archaeal and fungal OTU richness (alpha diversity) by performing a two-way analysis of variance (ANOVA, ɑ = 0.05), transforming the data for normality and homogeneity of variance when necessary. We then visualized similarities in microbial community composition between tree species and grove using non-metric multidimensional scaling (NMDS) of the Jaccard (presence-absence) and Bray-Curtis (relative abundance) dissimilarity metrics. To determine if beta diversity differed significantly between tree species and grove, we performed the multivariate permutation test perMANOVA using the ‘adonis’ function in the R VEGAN package (permutations = 999; Oksanen *et al.*, 2012). Because an assumption of perMANOVA is equal variance between groups, we also performed an analysis of multivariate homogeneity of group dispersions (permDISP; Supplemental Information Table S1).

In addition to running beta diversity analyses that included all taxa, we ran separate analyses for two subsets of fungal taxa that included only EMF or AMF. This allowed us to determine whether these root symbionts differed significantly between samples (by tree and grove). Specifically, EMF taxa were bioinformatically parsed as previously established (Glassman *et al.*, 2015) and AMF were bioinformatically parsed to include only individuals of Glomeromycotina. The resultant OTU tables were rarefied to even sampling depths (EMF = 3022; AMF = 40), visualized using NMDS, and analyzed using perMANOVA in the same way as described above.

As a complement to these multivariate tests, we compared the relative abundance of bacterial/archaeal and fungal phyla within each grove using non-parametric Mann-Whitney U test on ranks. Given that abundant taxa tend to contribute significantly to ecosystem functioning (Dai *et al.*, 2016), we also analyzed the frequency and relative abundance of dominant OTUs across species and grove. Specifically, we filtered both bacterial/archaeal and fungal OTU tables to include only those OTUs that comprised >1% of the total sequences. While 28 fungal OTUs met this criterion, only one bacterial/archaeal OTU met it (a *Bradyrhizobium* species); therefore, we summarized the results of the fungal OTUs only. Specifically, we visualized OTU frequency (presented as a percentage of the total number of samples) and relative abundance, and assessed significant differences in OTU relative abundance within each grove using non-parametric Mann-Whitney U tests.

Finally, we identified core OTUs that were unique to each tree species across both groves, where a core OTU was defined as an OTU that occurred in 100% of the samples recovered beneath a tree species. This enabled us to capture and identify the microbial members that were shared among all giant sequoia or sugar pine soils, but not both (i.e., the unique or host-specific core microbiome of each tree species). Core OTUs were identified using the compute_core_microbiome.py script in QIIME and graphically represented using Venn diagrams. Core bacterial/archaeal OTUs that were unique to each tree species were summarized at the phylum level. In addition, we determined the taxonomy of core bacterial/archaeal OTUs that were significantly more frequent (> 20% difference) beneath giant sequoia than sugar pine using EzTaxon (Kim *et al.*, 2012), and illustrated the presence/absence of these OTUs from all samples with the heatmap3 package (Zhao *et al.*, 2014). We provide no such summary for fungi, as neither giant sequoia nor sugar pine had a core microbiome based on our 100% occurrence definition.

In addition to determining differences in microbial community structure, we aimed to identify whether any structural differences may be attributed to tree-induced changes in soil parameters. To that end, univariate Spearman rank correlations were conducted to determine if any of the measured physiochemical parameters correlated with microbial richness. In addition, we used simple and partial Mantel tests to examine correlations between each physiochemical parameter and microbial community composition. A simple Mantel test is a non-parametric method that compares two distance matrices, and which calculates a correlation coefficient and p-value using permutations. It determines whether samples that are similar in one measure (e.g., soil pH) are similar in another (e.g., microbial composition). A partial Mantel test determines the relationship between two variables while holding the effects of other, potentially confounding, variables constant. We performed these Mantel tests using Jaccard measures of dissimilarity for microbial composition, and Euclidean distances for each soil parameter. Because we were particularly interested in understanding if and how giant sequoia influence microbial communities via changes in the soil, we only included physiochemical parameters that were found to differ significantly by tree species in our analyses (Hart et al., unpublished data), and we conducted separate analyses for each grove. Namely, we determined whether bacterial/archaeal and fungal community composition correlated with soil pH, extractable aluminum (Al), sum of the base cations (composed of 76-91% calcium, 5-16% magnesium, 2-11% potassium, and 0-3% sodium), and gravimetric soil moisture. All analyses were conducted in R version 3.2.1 (R Core Team, 2017).

## Results

### Characterizing Giant Sequoia Bacterial/Archaeal Communities (Q1)

Averaged across both the Mariposa and Merced groves, Proteobacteria comprised the majority of the 16S sequences recovered beneath giant sequoia (31.5% ± SE 0.5%), followed by Acidobacteria (15.9% ± SE 0.5%), Actinobacteria (15.6% ± SE 0.4%), Planctomycetes (11.2% ± SE 0.3%), and Verrucomicrobia (9.0% ± SE 0.2%; Supplemental Information Figure S1A). Together, these five phyla accounted for greater than 80% of the sequences. Forty-one less abundant phyla were also recovered from giant sequoia soils, three of which were archaeal.

### Characterizing Giant Sequoia Fungal Communities (Q1)

Averaged across both the Mariposa and Merced groves, Basidiomycota comprised the majority of ITS sequences recovered beneath giant sequoia (82.1% ± SE 2.4%), followed by Ascomycota (12.5% ± SE 1.5%), and Zygomycota (2.1% ± SE 1.4%). Less than 2% were Glomeromycota or unidentified fungi (Supplemental Information Figure S1B). *Hygrocybe* was the most dominant Basidiomycota genus recovered, comprising 21.5% (± SE 5.8%) of all sequences. *Wilcoxina* was the most relatively abundant Ascomycota genus recovered from beneath giant sequoia, comprising 4.9% (± SE 1.1%) of all sequences.

### Comparing giant sequoia and sugar pine microbial communities across both groves: Alpha and beta diversity (Q2 & Q3)

Bacterial/archaeal communities were most strongly structured by grove effects, followed by host tree differences (Figure 1A, Table 1A). Bacterial/archaeal richness was greater under giant sequoia (mean richness ± SE, Mariposa Grove: 5,924.6 ± 129.2, Merced Grove: 6,164.6 ± 104.6) compared to sugar pine (mean richness ± SE, Mariposa Grove: 5,327.7 ± 178.5, Merced Grove: 5,875.4 ± 221.5; Figure 2A). These differences remained constant across grove (no significant tree x grove interaction, P > 0.1). At the phylum level, Proteobacteria, Actinobacteria, and Gemmatimonadetes were relatively more abundant—and Acidobacteria, Armatimonadetes, and TM7 were relatively less abundant—in giant sequoia compared to sugar pine soils (Supplemental Information Figure S1A). However, these differences were not consistent across groves and some were only marginally significant (P = 0.05-0.10).

**Figure 1.**
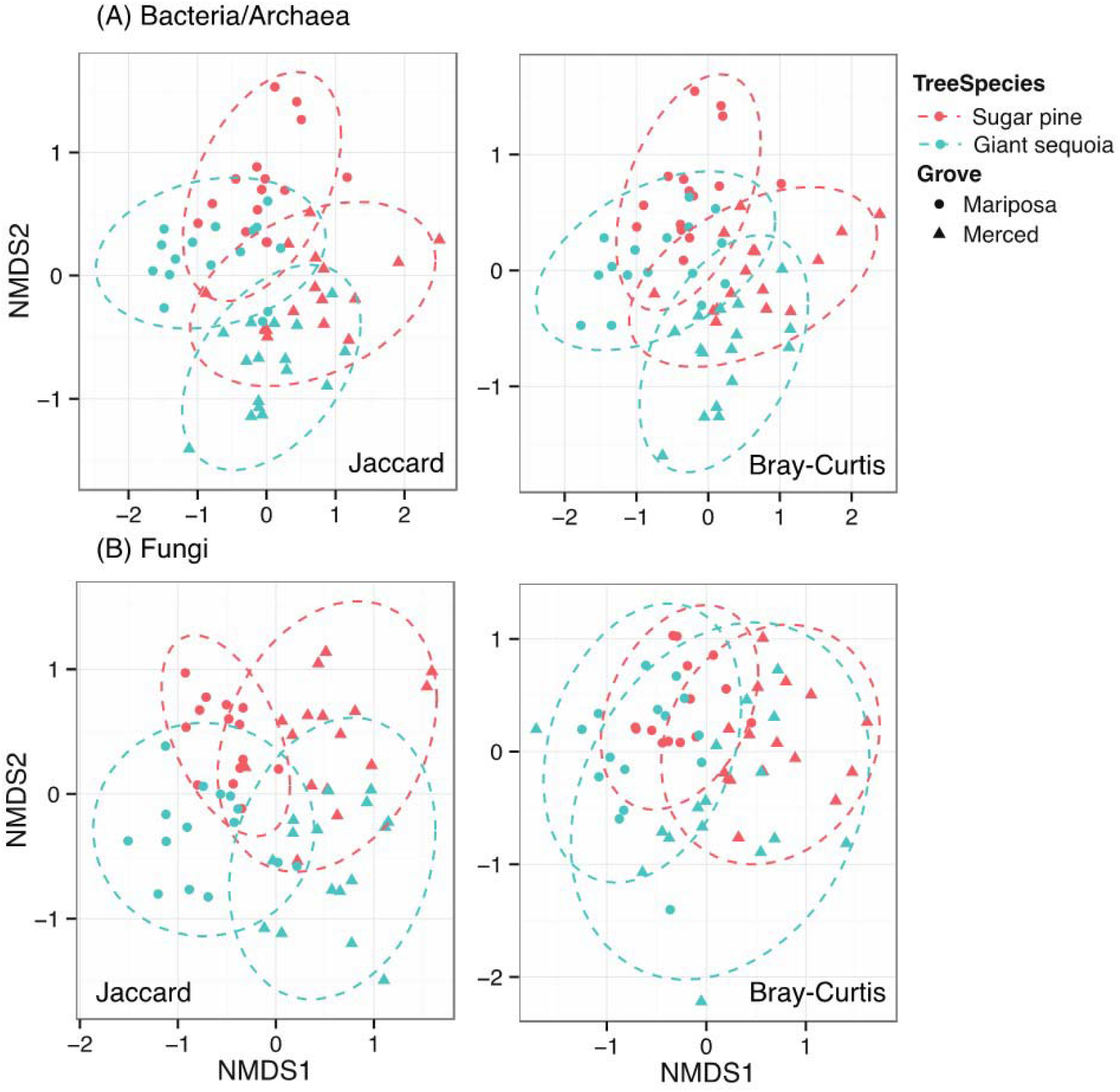
Influence of tree species (giant sequoia and sugar pine) and grove (Mariposa and Merced Grove) on (A) bacterial/archaeal and (B) fungal community composition. Left panel = non-metric multidimensional scaling (NMDS) of Jaccard (presence-absence) dissimilarity metric. Right panel = NMDS of Bray-Curtis (relative abundance) dissimilarity metric. Each symbol corresponds to a sample collected from one of two groves, and each color corresponds to a tree species. Points that are close together represent samples with similar community composition, and the dashed ovals represent 95% confidence intervals of sample ordination grouped by unique tree x grove combinations. The stress values for the bacterial/archaeal ordinations were 0.06 (Jaccard) and 0.07 (Bray-Curtis); the stress values for the fungal ordinations were 0.14 (Jaccard) and 0.18 (Bray-Curtis).

**Figure 2.**
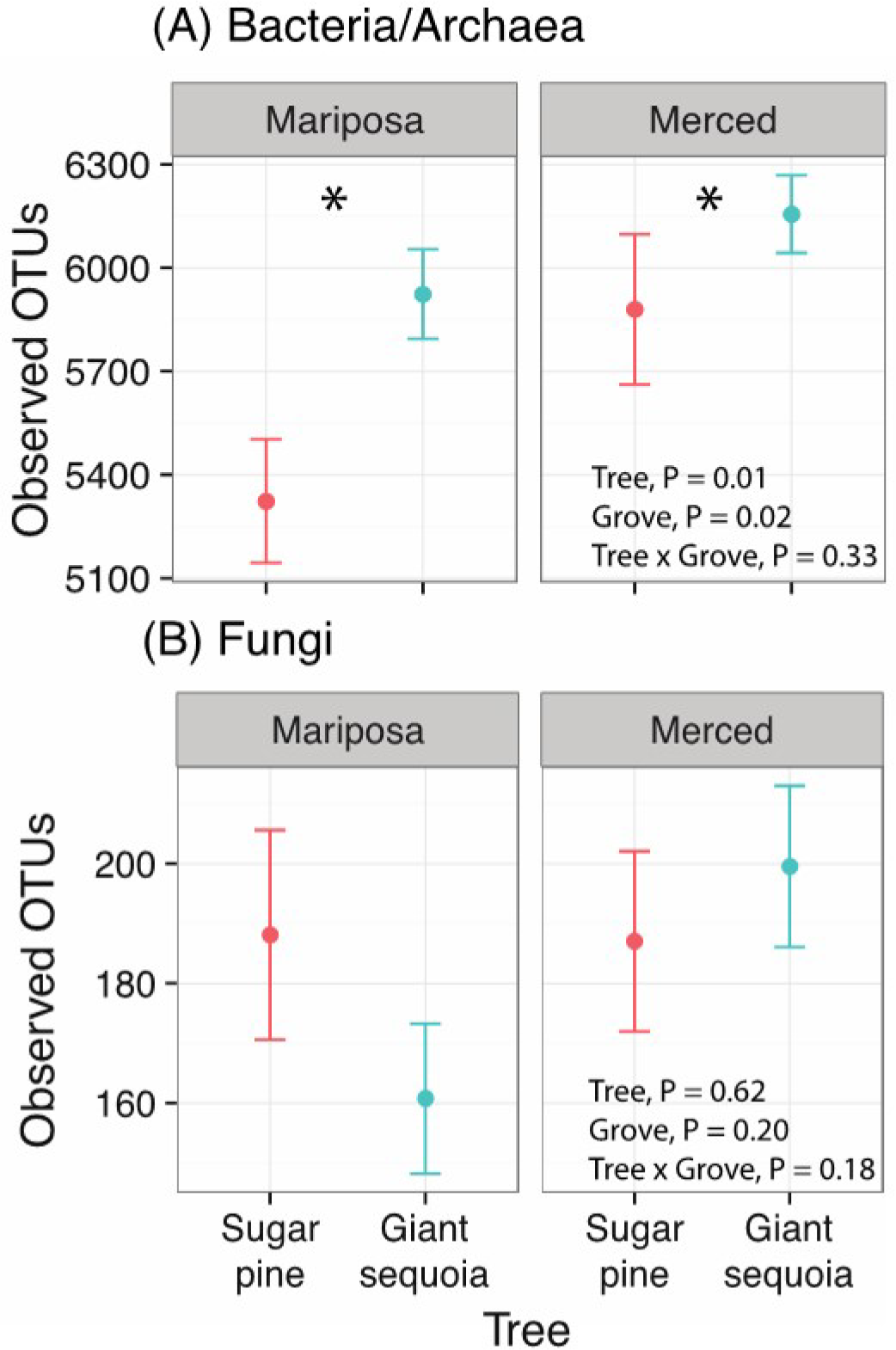
Influence of tree species (giant sequoia and sugar pine) and grove (Mariposa and Merced Grove) on alpha diversity for (A) bacteria/archaea and (B) fungi. Observed OTUs = number of observed operational taxonomic units, presented as mean ± 1 standard error (n =16). In (A), an asterisk denotes significant differences (P < 0.05) between tree species. P-values were derived from a two-way ANOVA.

**Table 1.** Results of the PERMANOVA for (A) bacteria/archaea and (B) fungi. Results are presented for both Jaccard and Bray-Curtis dissimilarity matrices. Because there was a significant tree x site interaction for fungi, we present the effects of tree species separated by grove. Site = Mariposa versus Merced Grove. Tree Species = giant sequoia versus sugar pine.

| **(A) Bacteria/Archaea** |  | **F<sub>model</sub>** | **R<sup>2</sup>** | **P value** |
| --- | --- | --- | --- | --- |
| *Jaccard* |  |  |  |  |
| Site |  | 3.663 | 0.057 | 0.001 |
| Tree Species |  | 2.29 | 0.036 | 0.001 |
| Site x Tree |  | 1.16 | 0.018 | 0.088 |
| *Bray-curtis* |  |  |  |  |
| Site |  | 9.427 | 0.134 | 0.001 |
| Tree Species |  | 4.09 | 0.063 | 0.001 |
| Site x Tree |  | 1.296 | 0.017 | 0.145 |
| **(B) Fungi** |  | **F<sub>model</sub>** | **R<sup>2</sup>** | **P value** |
| *Jaccard* |  |  |  |  |
| Site |  | 4.06 | 0.064 | 0.001 |
| Tree Species |  | 2.32 | 0.038 | 0.001 |
| Site x Tree |  | 1.34 | 0.021 | 0.016 |
|  | Tree Species in Merced Grove | 1.98 | 0.062 | 0.001 |
|  | Tree Species in Mariposa Grove | 1.83 | 0.064 | 0.002 |
| *Bray-curtis* |  |  |  |  |
| Site |  | 3.589 | 0.047 | 0.001 |
| Tree Species |  | 1.62 | 0.027 | 0.007 |
| Site x Tree |  | 1.38 | 0.022 | 0.032 |
|  | Tree Species in Merced Grove | 1.79 | 0.056 | 0.001 |
|  | Tree Species in Mariposa Grove | 1.26 | 0.045 | 0.096 |

Fungal communities were also structured most strongly by grove effects, followed by host tree differences (Figure 1B; Table 1B). However, in contrast to bacteria/archaea, there was a significant tree by grove interaction (P < 0.05), and fungal richness did not significantly differ between tree species (giant sequoia mean richness ± SE, Mariposa Grove: 160.7 ± 12.7, Merced Grove: 200.1 ± 13.1; sugar pine mean richness ± SE, Mariposa Grove: 187.9 ± 17.7, Merced Grove: 187.3 ± 14.8; Figure 2B). At the phylum level, Basidiomycota and Glomeromycota were relatively more abundant—and Ascomycota and Zygomycota were relatively less abundant—in giant sequoia compared to sugar pine soils (Supplemental Information Figure S1B). These phylum-level differences were not consistent across groves and some were marginally significant (P = 0.05-0.1). In addition, the composition of EMF and AMF communities showed small but significant effects of tree host and grove with no interaction (Supplemental Information Figure S2)—and AMF had significantly more OTUs beneath giant sequoia (giant sequoia mean richness ± SE = 8.5 ± 1.09; sugar pine mean richness = 5.4 ± 0.96; P = 0.009).

### Comparing giant sequoia and sugar pine microbial communities across both groves: Dominant taxa and core members (Q2 & Q3)

Only one bacterial OTU, a *Bradyrhizobium* (phylum Proteobacteria), comprised more than 1% of the total bacterial sequences. In contrast, 28 fungal taxa each comprised more than 1% of the total fungal taxa (Figure 3). Of these 28 fungal taxa, 64% were EMF and 18% belong to the genus *Hygrocybe*. We observed a number of these dominant OTUs whose frequency, relative abundance, or both differed consistently between tree species. For example, in both Mariposa and Merced groves, an unidentified *Cryptococcus* species was recovered from 100% of samples, but was relatively more abundant beneath sugar pine. An unidentified *Byssocorticium* species, an EMF taxon, also showed a consistent trend across groves, where it was more frequent and relatively abundant beneath sugar pine compared to giant sequoia. In contrast, species of *Hygrocybe*, which is generally considered saprophytic (although see discussion below), were almost always more frequent and relatively more abundant in giant sequoia soils, although these differences were not always statistically significant (Figure 3). Finally, some OTUs differed significantly between tree species in one grove but not the other. This included *Russula acrifolia*, another EMF taxon (Tedersoo *et al.*, 2010), which was more frequent and relatively abundant beneath sugar pine than giant sequoia in the Mariposa grove, and an unidentified *Geminibasidium* species, which is a xerotolerant basidiomycete yeast (Nguyen *et al.*, 2013), was relatively more abundant beneath sugar pine than giant sequoia in the Merced grove.

**Figure 3.**
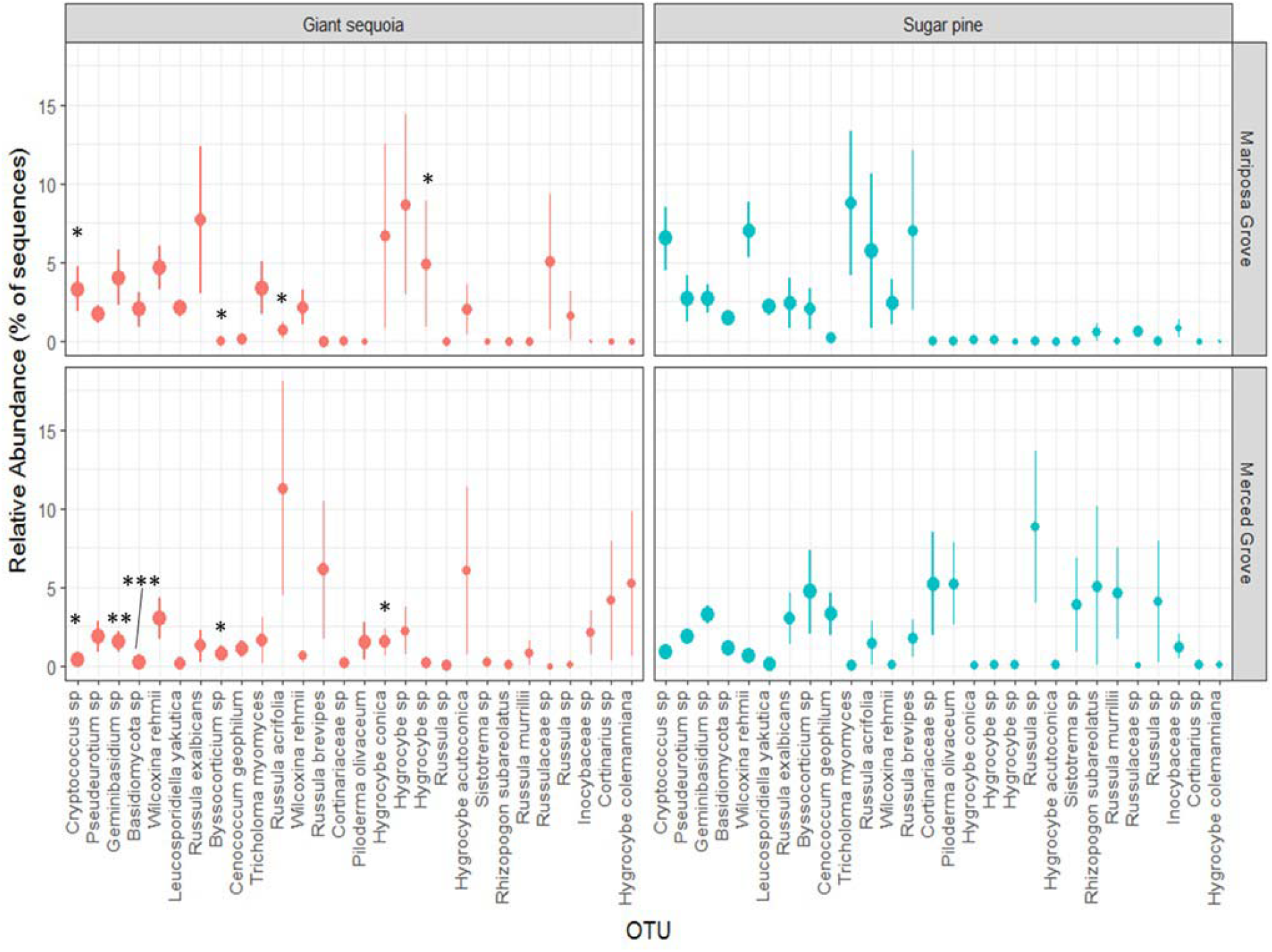
The relative abundance of the most abundant fungal OTUs in giant sequoia and sugar pine soils across both groves. Error bars = 1 standard error of the mean (n = 16). The size of each point is scaled by the frequency of an OTU (how many samples it was recovered from), with larger circles corresponding to greater frequency. Significant differences in OTU relative abundance between tree species were assessed using Mann-Whitney U test on ranks (* p < 0.05, ** p < 0.01, *** p < 0.001).

There were very few fungal OTUs that were recovered from 100% of giant sequoia or sugar pine samples in either grove. Accordingly, neither giant sequoia nor sugar pine had a core fungal microbiome comprised of OTUs that were recovered from all samples (Figure 4A). Even when we relaxed the definition of a core to require only 80% frequency, giant sequoia soils contained only two unique core fungal OTUs and sugar pine soils contained one—both of which were Zygomycota. In contrast, bacterial/archaeal communities beneath giant sequoia and sugar pine harbored a number of OTUs that made up a unique core (i.e., were recovered from 100% of giant sequoia samples or 100% of sugar pine samples, but not 100% of both; Figure 4B). Both tree species contained the phyla Proteobacteria, Acidobacteria, Planctomycetes, Actinobacteria, Bacteroidetes, and Verrucomicrobia in their core community (Figure 4C & D). Each core also contained a number of OTUs representing phyla unique to that tree species. For instance, giant sequoia contained core members from Chloroflexi, Gemmatimonadetes, and TM7, while sugar pine did not (Figure 4C). Similarly, only sugar pine contained core members from Armatimonadetes, Chlorobi, and OD1 (Figure 4D). In addition to these phylum-level differences, the size of the core for giant sequoia and sugar pine differed, with giant sequoia containing substantially more core OTUs (101 OTUs) than sugar pine (50 OTUs; Figure 4B). However, of the 101 core OTUs beneath giant sequoia, only 13% were notably less frequent (frequency < 80%) in sugar pine soils (Figure 4E; Supplemental Information Table S2). Similarly, of the 50 core OTUs beneath sugar pine, only 10% were notably less frequent in giant sequoia soils (Supplemental Information Table S2). In all other cases, OTUs that comprised the core of one tree community were often missing from only a few samples in the other tree community (80-95% frequency).

**Figure 4.**
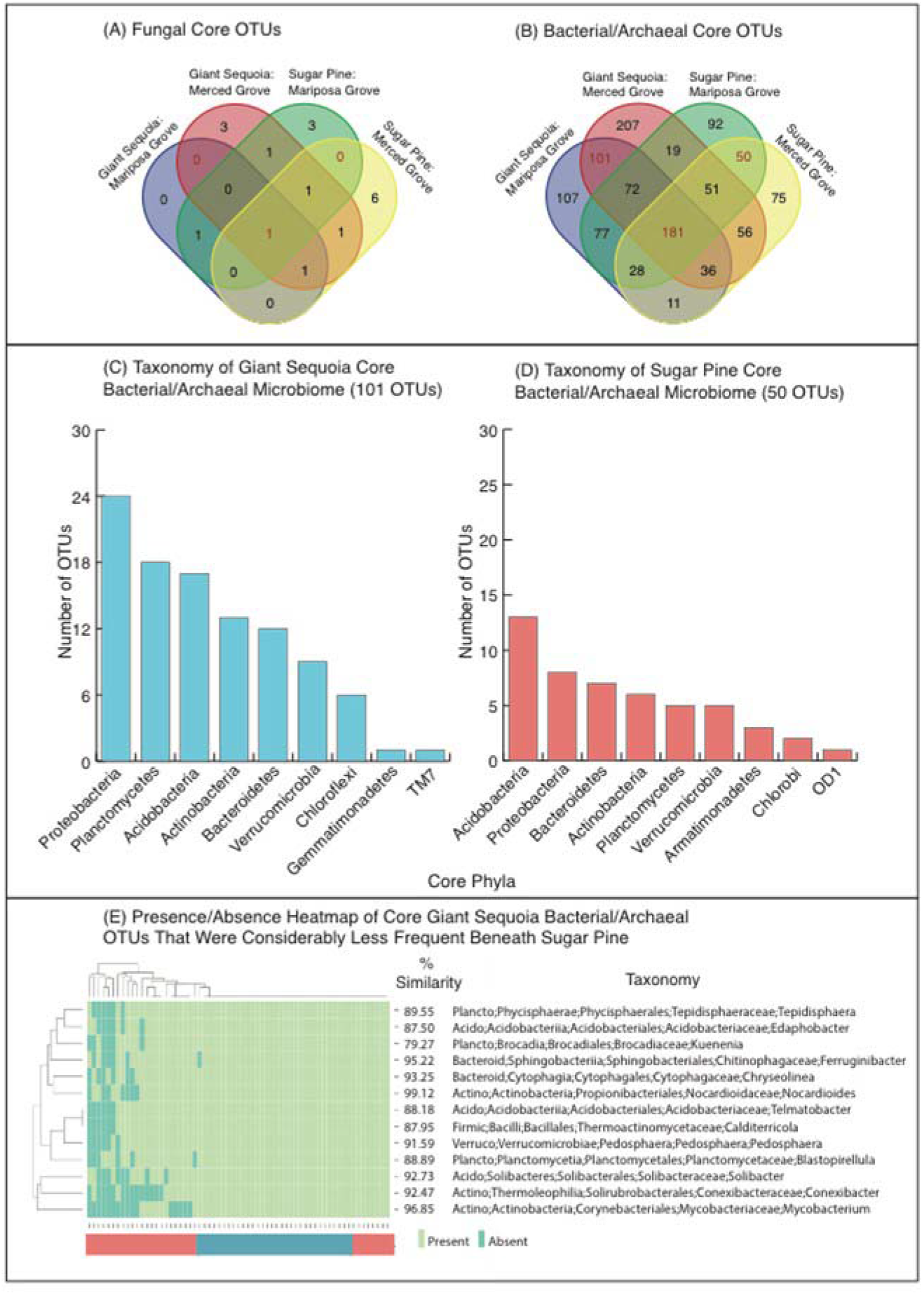
Shared OTUs of the (A) fungal and (B) bacterial/archaeal core microbiomes associated with giant sequoia and sugar pine. The Venn diagrams show absolute number of OTUs shared between core microbiomes of each tree species across two groves. Phylum level taxonomic information is also provided for the OTUs comprising the (C) giant sequoia and (D) sugar pine core bacterial/archaeal communities. (E) Heatmap illustrating presence/absence and taxonomy of giant sequoia’s core OTUs that were considerably less frequent (20% difference) in sugar pine soils. Taxonomic information was derived from EzTaxon, and % similarity is the sequence similarity between the OTU and its nearest cultured match. Colors of the bar beneath the heatmap correspond to tree type (pink = sugar pine, blue = giant sequoia).

### Determining the indirect effects of tree species on microbial diversity and composition (Q4)

Giant sequoia and sugar pine soils differed significantly in a number of measured physiochemical parameters (Supplemental Information Table S3). Of these parameters, bacterial/archaeal and fungal community composition correlated most strongly and consistently with soil pH (Figure 5); these relationships with soil pH remained strong even when the effects of other soil parameters were statistically controlled for using a partial Mantel test. The composition of bacterial/archaeal and fungal communities also tended to be strongly related to differences in soil moisture (gravimetric water content) in both the simple and partial Mantel tests. In addition, bacterial/archaeal and fungal community composition correlated with extractable aluminum and the sum of base cations; however, these relationships were inconsistent between groves for fungi, and often disappeared for both microbial groups when the effects of other soil parameters were accounted for (Figure 5).

**Figure 5.**
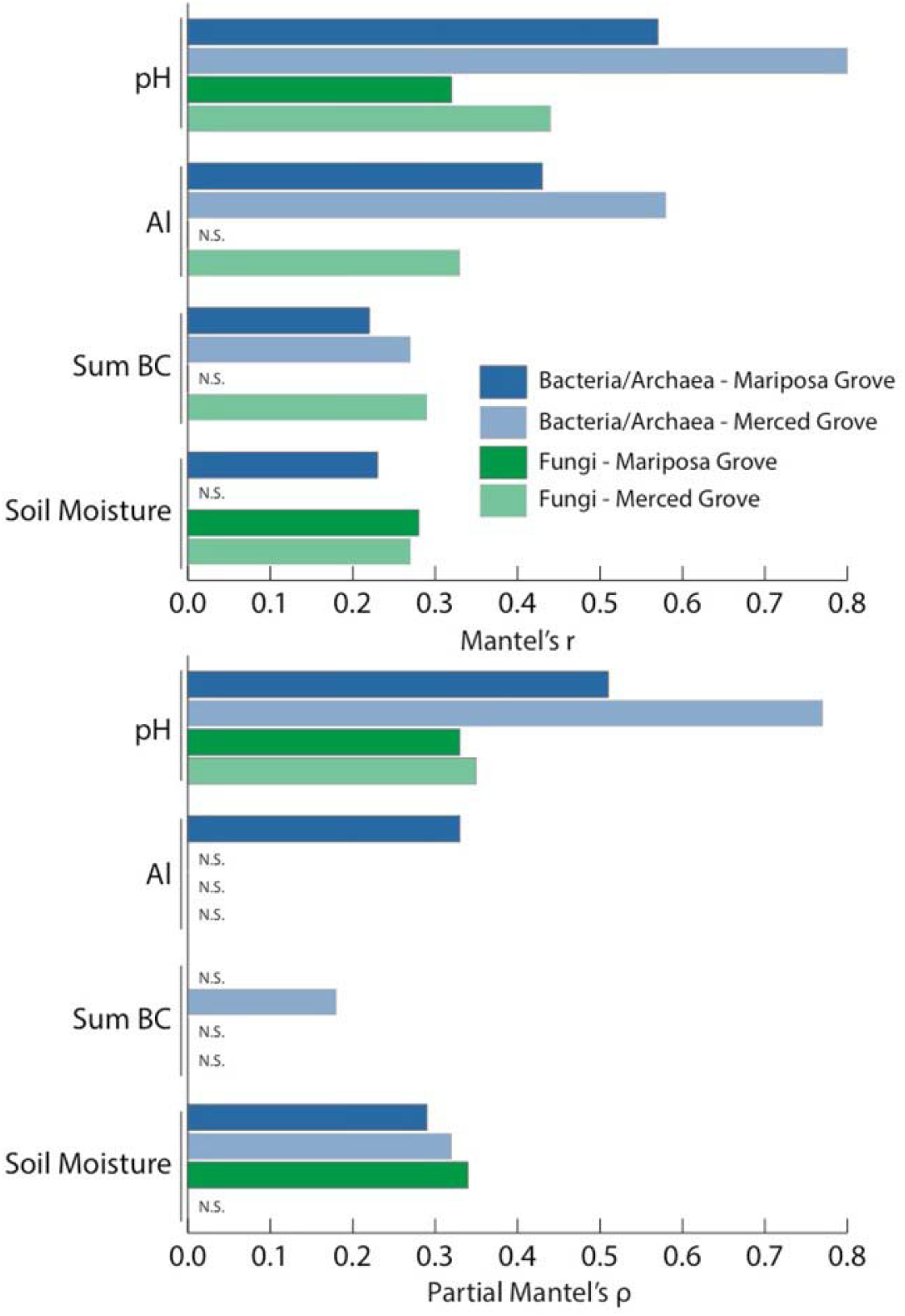
Results of (top panel) simple (r) and (bottom panel) partial (ρ) Mantel tests relating soil parameters that varied between tree types to bacterial/archaeal and fungal community composition. Only those parameters that correlated significantly with composition at least once are presented. pH = soil pH; Al = soil xtractable aluminum; Sum BC = sum of the base cations (Ca^2+^, Mg^2+^, K^+^, and Na^+^); soil moisture = soil gravimetric water content. N.S. = non-significant (p > 0.05) correlation.

In contrast to microbial composition, bacterial/archaeal and fungal richness were largely unrelated to the measured physiochemical parameters (Supplemental Information Table S3). Only in the Merced Grove did bacterial/archaeal richness correlate positively with pH, and negatively with soil moisture and extractable aluminum. Similarly, of all the physiochemical parameters, fungal richness only correlated (positively) with bulk density in the Mariposa Grove, and this relationship was only marginally significant (P = 0.05-0.1).

## Discussion

Giant sequoia are the largest, and some of the longest-lived, trees in the world. An iconic species with great ecological and cultural importance, little is known about how giant sequoia individuals interact with belowground communities. In this study, we used next-generation sequencing techniques to describe for the first time the soil microbiome associated with this charismatic megaflora. We compared giant sequoia microbiomes to those of co-occurring sugar pine trees across two groves on contrasting geological substrates in Yosemite National Park, USA, and determined whether host tree differences in microbial communities were related to differences in soil physiochemical parameters. While this kind of observational study design—which is largely unavoidable given the long-lived nature of the host trees under study—limits our power of causal inference, these limitations can be partly addressed by future work that tracks giant sequoia soil microbiomes from time since establishment and across additional groves with (dis)similar parent materials.

### Characterizing giant sequoia soil microbiomes and comparing them to sugar pine microbiomes across both groves (Q1 - Q3)

Differences in bacterial/archaeal community composition between groves tended to be greater than those differences associated with tree species. Still, there was evidence that giant sequoia influenced underlying bacteria and archaea in unique ways compared to sugar pine. Specifically, we found that bacterial and archaeal richness was greater beneath giant sequoia than sugar pine (Figure 2A), possibly because giant sequoia are larger and create more niche space within the soil (Glassman *et al.*, 2017). In addition, communities of bacteria/archaea were compositionally distinct from those beneath sugar pine (Figure 1A). These differences remained constant across the two groves, despite the fact that soils from each grove were derived from geochemically distinct substrates. Fungal community composition also differed between giant sequoia and sugar pine; however, the specific ways that fungi differed between tree species depended on the grove (Figure 1B). Overall, these findings—which agree with a number of other studies where bulk soil microbial communities differed between contrasting tree species (Ayres *et al.*, 2009; Thoms *et al.*, 2010; Scheibe *et al.*, 2015; Sun *et al.*, 2016)—suggest that, despite relatively large grove differences, a “host signal” can still be observed.

Giant sequoia and sugar pine are known to associate with two contrasting groups of mycorrhizal fungi. The former with AMF (Fahey *et al.*, 2012) and the latter with EMF (Walker, 2001). While AMF and EMF were recovered from beneath both tree hosts— possibly because of understory influences or root overlap—communities associated with giant sequoia differed from those associated with sugar pine (Supplemental Information Figure S2). AMF communities were considerably more diverse beneath giant sequoia, and EMF taxa such as an unidentified *Byssocorticium* species and *Russula acrifolia* were more frequent and relatively abundant beneath sugar pine. While this is to be expected, our findings show that differences in mycorrhizal communities in mixed stands of AMF and EMF trees can be seen at the bulk soil scale, differences that can have cascading effects on global scale biogeochemical processes including the cycling of carbon (Averill *et al.*, 2018) and nitrogen (Mushinski *et al.*, 2019).

Microbial taxa that are abundant within the community can contribute significantly to ecosystem function (Dai *et al.*, 2016). To discern patterns in frequent and abundant taxa between trees, we filtered both the bacterial/archaeal and fungal OTU tables to include only those OTUs that comprised greater than 1% of the total sequences. In accordance with the idea that prokaryotic microbial communities often include a very small number of dominant members, and are instead comprised of many rare members (Sogin *et al.*, 2006), only one bacterium passed our high abundance filter. This bacterial OTU, which was of the genus *Bradyrhizobium*, comprised 2.1% of the total 16S rRNA sequences recovered, making it the dominant bacterial OTU at both groves. Recent work employing quantitative population genomics suggests that free-living *Bradyrhozbia* are unexpectedly incapable of fixing atmospheric N and may instead metabolize aromatic carbon sources (VanInsberghe *et al.*, 2015), which may explain why this genus tends to dominate microbial communities of forest bulk soil (Uroz *et al.*, 2010; Hartmann *et al.*, 2012; VanInsberghe *et al.*, 2015).

In contrast to bacteria/archaea, we recovered 28 abundant OTUs that each comprised greater than 1% of the total ITS sequences (Figure 3). Notably, 18% of these OTUs were from the genus *Hygrocybe* (waxcaps)*. Hygrocybe* are widespread and can be found in a variety of habitats worldwide (Halbwachs *et al.*, 2013a), including forests of the *Sequoiadendron* sister genus *Sequoia* (Enright *et al.*, 2019). For example, *Hygrocybe* was the dominant taxon comprising 13% of ITS1 Illumina MiSeq sequences in a California coastal redwood (*Sequoia sempervirens*) forest (Enright *et al.*, 2019). These fungi are frequently considered to be saprotrophic, although recent work has implicated some *Hygrocybe* species, such as *H. virginea*, as being endophytic (Halbwachs *et al.*, 2013b; Persoh, 2013; Tello *et al.*, 2014). In our study, *Hygrocybe* were rarely recovered from beneath sugar pine, indicating a preference for giant sequoia soils. Indeed, a canonical discriminant analysis (CDA) illustrated that differences in *Hygrocybe* drove much of the fungal compositional variability between the two trees (data not shown). While the exact trophic lifestyle of this genus remains uncertain (Seitzman *et al.*, 2010), it is possible that—given the preponderance of *Hygrocybe* species in giant sequoia soils and their apparent ability to act as root and systemic endophytes—these fungi may form ecologically significant symbiotic relationships with giant sequoia (Zarraonaindia *et al.*, 2015). Future research should therefore identify whether such a relationship exists and, if so, what the ecological implications may be for giant sequoia growth and survival.

In addition to identifying common and abundant OTUs, it can be useful to distinguish core members of a microbial community that remain constant across space or time (Shade & Handelsman, 2012). Doing so helps define a healthy (or alternatively a degraded) community, and can improve our understanding how that community will respond to future perturbations. In contrast to fungi, which had no discernable core community, we identified considerable core bacterial/archaeal communities associated with both tree species (Figure 4A & B). Interestingly, giant sequoia’s unique core was double the size of sugar pine’s, indicating that this giant, long-lived tree maintains a relatively large and consistent set of prokaryotic OTUs in its surrounding soil. The larger community associated with giant sequoia could be due to its age and size, as larger trees are known to host more microbial taxa (Glassman *et al.*, 2017). Of these OTUs, thirteen were also considerably (at least 20%) less frequent in sugar pine soils. However, there were no clear trends in the taxonomy or ecology of these thirteen OTUs, with family associations ranging from *Norcardioidaceae* (contains endophytes and species capable of degrading organic matter; Tóth & Borsodi, 2014) to *Mycobacteriaceae* (contains animal pathogens and species capable of degrading hydrocarbons; Lory, 2014). Metagenomic data could provide a more complete picture of core community dynamics, as some evidence suggests that communities assemble at the functional rather than the phylogenetic level (Burke *et al.*, 2011). Regardless, our data contribute to a small set of previously published work that explicitly identify core communities in bulk soil (Andrew *et al.*, 2012; Orgiazzi *et al.*, 2013), and indicate that giant sequoia and sugar pine each harbor unique core communities of bacteria/archaea, but lack consistent core fungal OTUs at the spatial scale studied here.

It is possible that having a large and diverse core community of bacteria/archaea aids in the long-term success of giant sequoia individuals. In addition to providing critical biogeochemical functions (e.g., decomposition of organic matter and nutrient cycling), core and abundant microorganisms within soil may act as a source that “seeds” rhizospheric and endospheric communities, ultimately contributing directly to plant health. Indeed, it has been proposed that some foliar endophytes persist through the winter as saprobes of litter only to re-invade host leaves in the spring (Unterseher *et al.*, 2013; Baldrian, 2017). A recent study assessing foliar bacterial endophytic communities of giant sequoia and coastal redwoods found that giant sequoia contained a diverse endophytic community, with major phyla including Acidobacteria, Actinobacteria, Bacteroidetes, Firmicutes, Fusobacteria, Proteobacteria, and TM7 (Carrell & Frank, 2015). Notably, five of the twenty dominant orders recovered from giant sequoia foliage samples were represented in our core bacterial/archaeal community dataset (Actinomycetales, Burkholderiales, Rhizobiales, Rhodospirillales, and Sphingobacteriales). It is conceivable that at least some of these endophytes are derived from the soil community; however, (at minimum) comparative sampling of giant sequoia microbial communities within the same site, and (at maximum) use of more advanced tracing techniques (e.g., isotopes or quantum dots), will be required to assess this. Future studies that focus on connecting the giant sequoia holobiont with soil communities should provide promising insights to the stability of this tree species over millennia.

### Determining the indirect effects of tree species on microbial diversity and composition (Q4)

Trees can affect the composition and diversity of soil microbial communities through a variety of indirect mechanisms. Some of these mechanisms include changes in root exudation, nutrient uptake, soil microclimate, pH, and the amount and quality of litter inputs (Binkley & Giardina, 1998). In our study, a number of notable soil parameters differed between giant sequoia and sugar pine (Hart et al., unpublished data; Supplemental Information Methods S1)—and while bacterial/archaeal and fungal richness were generally insensitive to these parameters, community composition correlated with a number of them. Most strongly and consistently, bacterial/archaeal and fungal community composition related to soil pH and soil moisture (Figure 5). Soil moisture dynamics are coupled to water stress, oxygen diffusion, and substrate supply; differences in soil moisture can therefore alter metabolic activity of microbial functional groups, the occurrence of aerobic/anaerobic processes (e.g., aerobic decomposition/denitrification), and ultimately microbial community composition (Manzoni *et al.*, 2012). Soil pH is a well-known driver of bacterial/archaeal community composition and diversity, a driver that is often unrivaled by other soil parameters due to the narrow tolerance range of prokaryotic microorganisms (Lauber *et al.*, 2009; Collins *et al.*, 2016). While fungi are thought to exhibit wider pH ranges for optimal growth (Rousk *et al.*, 2010), a recent study indicates that soil pH can also be a strong mediator of fungal community composition—a finding consistent with ours (Glassman *et al.*, 2017).

There is growing recognition that calcium availability also regulates soil microbial (Grayston *et al.*, 2004; Allison *et al.*, 2007; Sun *et al.*, 2016; Glassman *et al.*, 2017) and faunal (Reich *et al.*, 2005) community composition, organic matter decomposition (Hobbie *et al.*, 2006; Hobbie *et al.*, 2007), and overall forest health (Battles *et al.*, 2014). Calcium provides structure to many cell walls, and the cycling of this cation in soil depends largely on decomposition (Chapin *et al.*, 2011). In our study, giant sequoia litter was significantly enriched in base cations compared to sugar pine, and this resulted in increased availability of exchangeable calcium (in addition to Mg^2+^, K^+^, Na^+^) in the upper mineral soil. Accordingly, Mantel tests revealed that bacterial/archaeal and fungal community composition correlated significantly with the sum of the base cations, a composite measure that was composed primarily of calcium. However, this relationship almost always disappeared when the effects of the other variables were accounted for (Figure 5), suggesting that the influence of calcium was mediated by other soil parameters—namely soil pH. Base cations compete with H^+^ and Al^3+^ for exchange sites on soil particle surfaces; therefore, the concentration of calcium in soil mediates soil pH, such that higher amounts of exchangeable calcium (and other base cations) create less acidic soils (Reich *et al.*, 2005). The idea that soil calcium indirectly mediates microbial composition via soil pH agrees with the conclusions of some other studies (Grayston *et al.*, 2004; Reich *et al.*, 2005); however, Allison *et al.* (2007) suggested that calcium may influence the soil community primarily via calcium effects on soil aggregation. Taken together, these data indicate that—by maintaining high soil pH, nutrient availability (i.e., calcium and other base cations), and moisture compared to sugar pine—giant sequoia indirectly influence the composition of underlying microbial communities.

### Conclusion

Using next-generation sequencing techniques, we show for the first time that microbial communities of bulk soil differ between giant sequoia and a co-occurring conifer, sugar pine. Namely, giant sequoia supported unique bacterial/archaeal and fungal community composition, greater bacterial/archaeal richness, and a large unique core community of bacteria/archaea. These host tree differences, which were at least partially driven by soil pH, calcium availability, and soil moisture, were discernible despite concurrently large grove effects. In some cases, the influence of host tree differed between the two groves under study, which were close in proximity but had contrasting parent material. Our findings suggest that the effects of giant sequoia extend beyond mycorrhizal mutualists to include the broader community at the bulk soil scale.

## Acknowledgements

The authors declare that they have no conflict of interest. We thank Teresa L. Fukuda for field and laboratory assistance and Judy Chung for help with ITS1 sequencing. We also thank the National Park Service for allowing us to sample soils within Yosemite National Park. This research was supported by grants from the NSF Research Experience for Undergraduates (DBI-1263407) and the Critical Zone Observatory Programs (EAR-1331939), as well as a USDA National Institute of Food and Agriculture Seed grant (CA-R-PPA-5101-CG) and RSAP grant (CA-R-PPA-5093-H).

