## Supplemental Information for "Soil microbial communities associated with giant sequoia: How does the world’s largest tree affect some of the world’s smallest organisms?"

**Supplemental Methods S1. Physicochemical Analyses**

Gravimetric water content was determined by oven-drying a 15-mL subsample of sieved soil for 48 h at 105 °C. Soil pH was measured using a pH electrode (Orion DUAL STAR meter, Thermo scientific, Waltham, MA, USA) after allowing 5 g of air-dry soil to equilibrate with 0.01 M CaCl_2_ solution (1:5 w/v) for 30 minutes. Ammonium and NO_3_^-^ concentrations were determined by extracting 5 g of field-moist soil with 25 mL of 2 M potassium chloride (KCl), filtered through Whatman No. 1 filter paper (preleached with deionized water), followed by flow injection analysis (QuikChem 8500, Lachat Instruments, Hach Company, Loveland, CO). Ammonium and NO_3_^–^ were determined by colorimetric methods on separate manifolds, the former using phenolate and the latter using a cadmium-copper reduction column. To determine anaerobically mineralizable N, at the time of KCl-extraction, an additional 15-mL subsample was placed in a 120 mL specimen cup and 50 mL of deionized water was added. The suspension was mixed gently and then incubated at 40 °C for 7 days. After this period, 50 mL of 4 M KCl were added and the sample was shaken, filtered and analyzed as above. Anaerobically mineralizable N was calculated as the difference in NH_4_^+^ concentrations between incubated and the initial unincubated samples. Available P was measured on air-dried soil using a Bray-1 extraction. Twenty-five mL of a mixed, 0.03 M NH_4_F + 0.025 M HCl solution to ≈2.5 g of air-dried soil in a 50 mL polyethylene centrifuge tube and shaken for 5 min at 150 cycles per minute. The suspension was then filtered through a quantitative filter paper and stored at < 4 °C until analysis using an ascorbic acid colorimetric method. Total Kjeldahl N and P were determined using the Kjeldahl digestion protocol on air-dry, finely ground subsamples. After digestion, samples were analyzed by flow-injection colorimetry using salicylate and molybdate-ascorbic acid methods, respectively. Total N, along with total C, δ^13^C, and δ^15^N, were also determined by EA-IRMS at UC Davis’ Stable Isotope Facility. Extractable elements (Na, K, Mg, Ca, Al, Fe, and S) in soil were measured adding 25.0 mL of 1 M NH_4_Cl to ≈2.5 g of air-dried soil in a 50 mL polyethylene centrifuge tube. The suspension was shaken for 30 min at 150 cycles per minute and then filtered through a quantitative filter paper and store at < 4 °C until analysis. Solutions were diluted 1:10 (v/v) with deionized water containing 1% HNO_3_ prior to analysis by ICP-AES.

**Supplemental Figure S1.** Relative abundances of (A) bacterial and archaeal phyla and (B) fungal phyla averaged across replicates (n = 16) from beneath giant sequoia and sugar pine individuals in Mariposa and Merced groves. Both graph and legend share the same order, sequential from bottom to top. For bacteria/archaea, ‘other’ indicates the combined relative abundance of 26 lesser abundant phyla. Significant (P < 0.05) and marginally significant (P = 0.05-0.1) differences in phyla between tree species are illustrated for each grove (derived from Mann-Whitney U test on ranks).

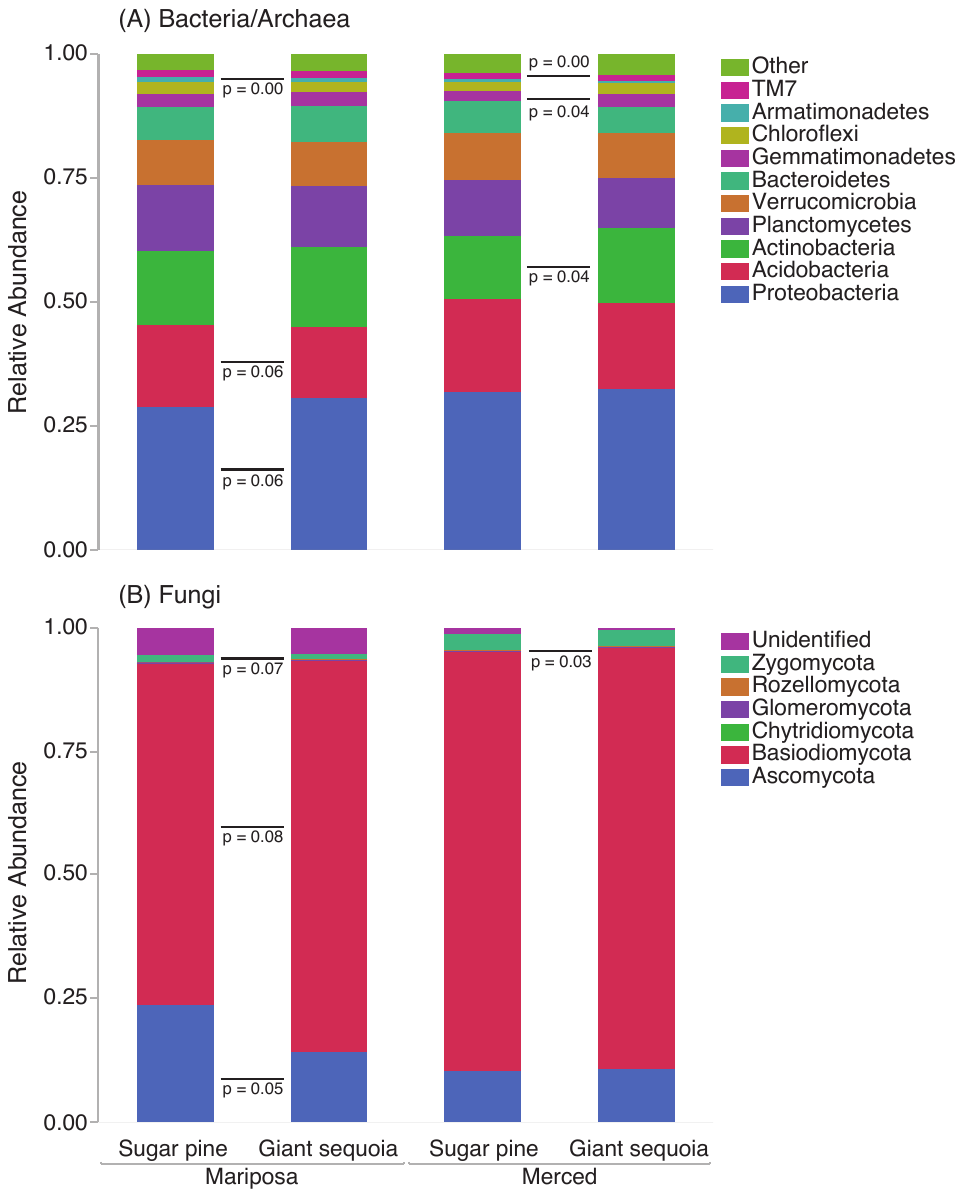

**Supplemental Figure S2.** Influence of tree species (giant sequoia and sugar pine) and grove (Mariposa and Merced Grove) on (A) ectomycorrhizal and (B) arbuscular mycorrhizal community composition. Figures are non-metric multidimensional scaling (NMDS) of Jaccard (presence-absence) dissimilarity metric. Each color corresponds to a sample collected from a particular tree x grove combination. Points that are close together represent samples with similar community composition. perMANOVA and PERMDISP results also shown from Jaccard dissimilarity metric for both (A) and (B). The stress values for (A) ectomycorrhizal fungi and (B) arbuscular mycorrhizal fungi were 0.20 and 0.18, respectively.

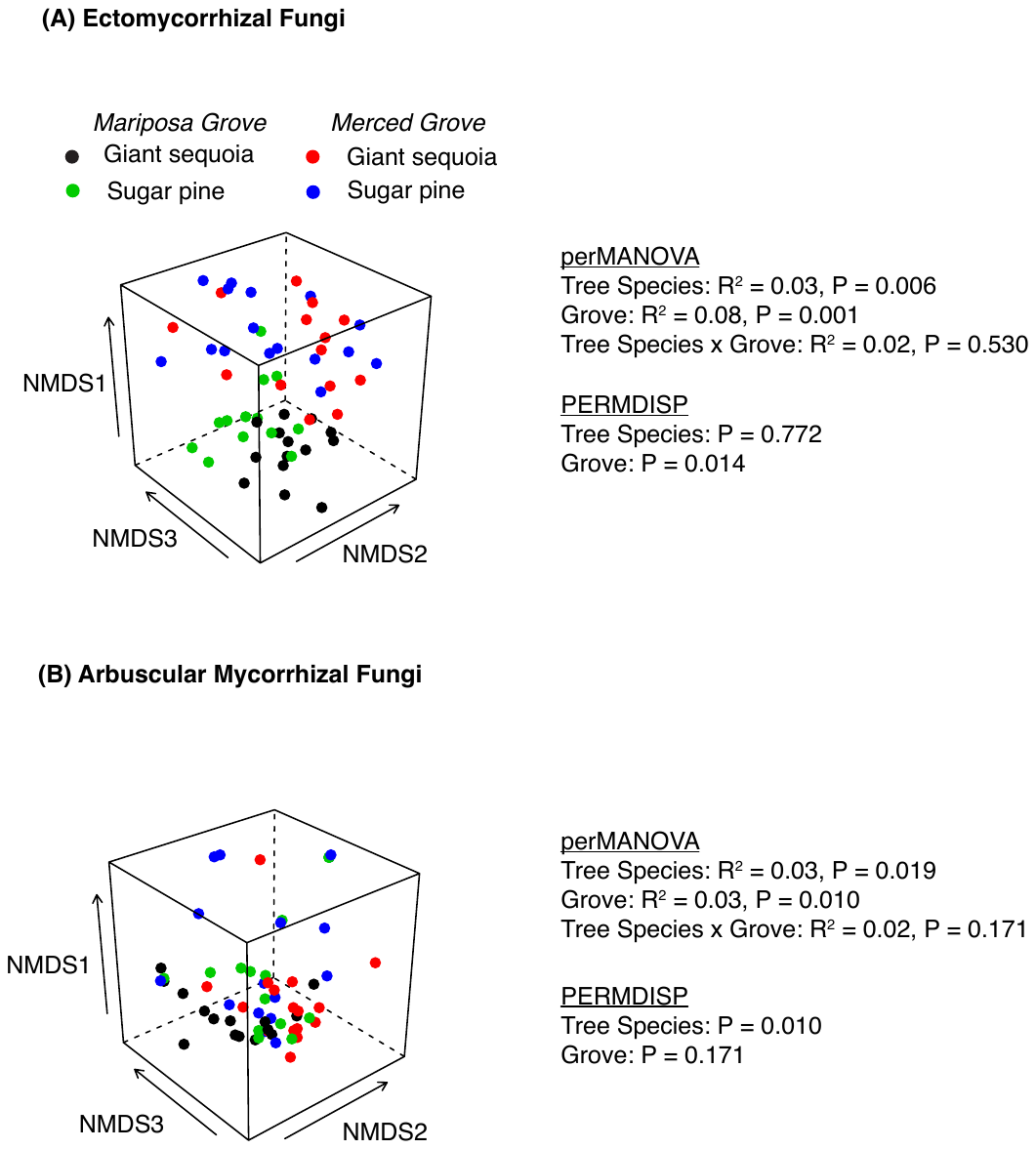

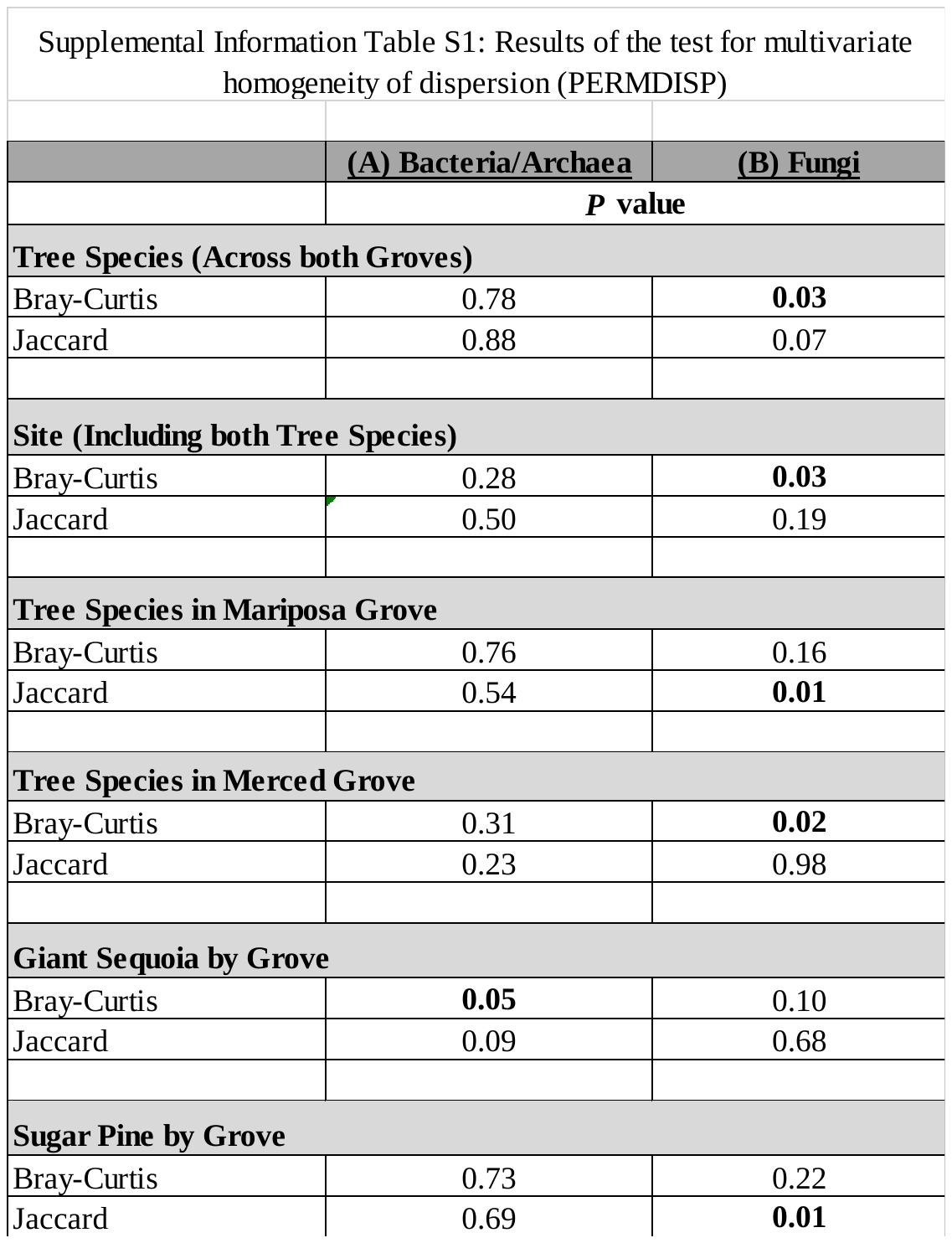

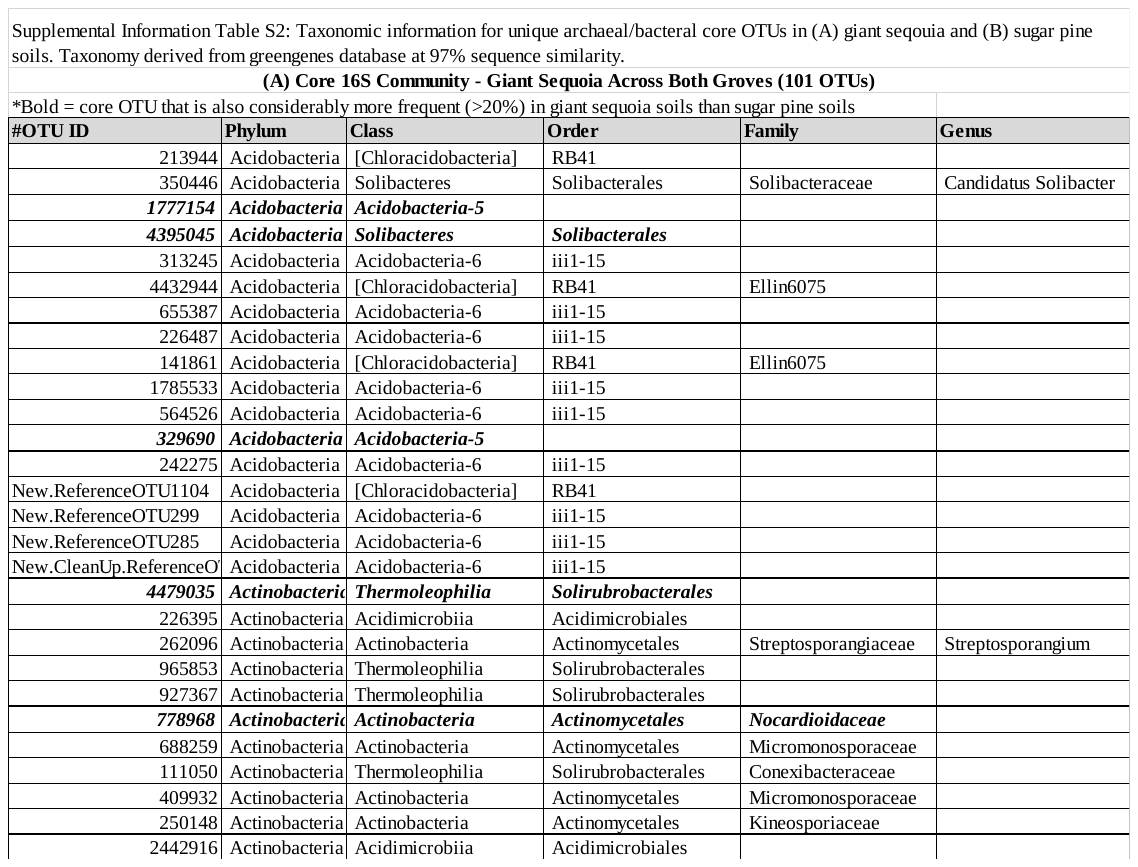

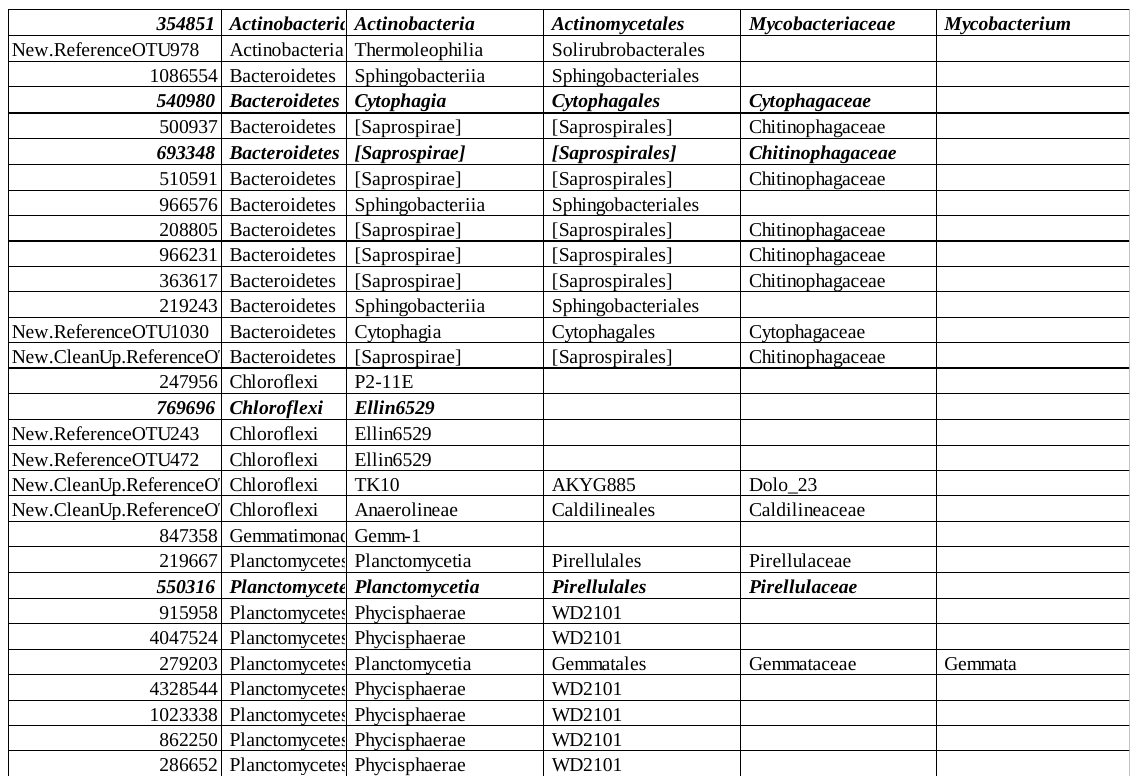

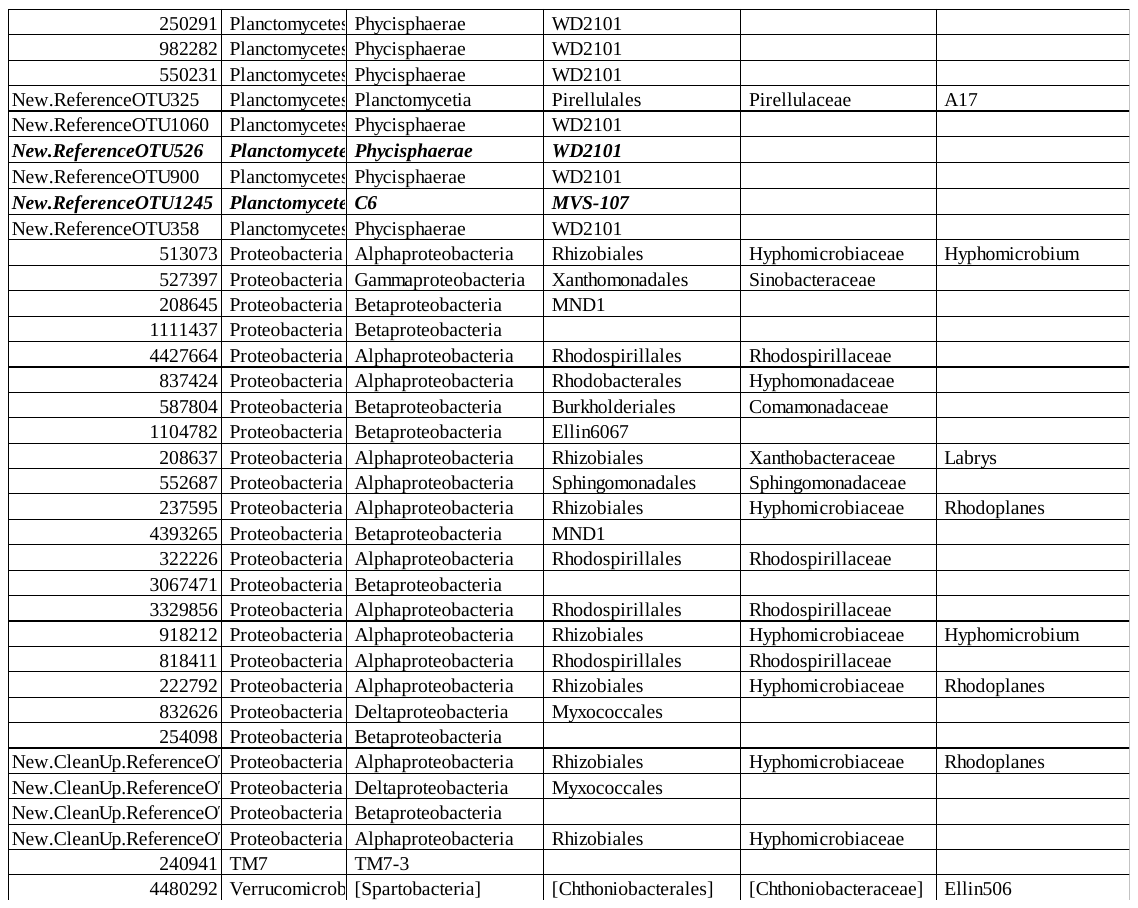

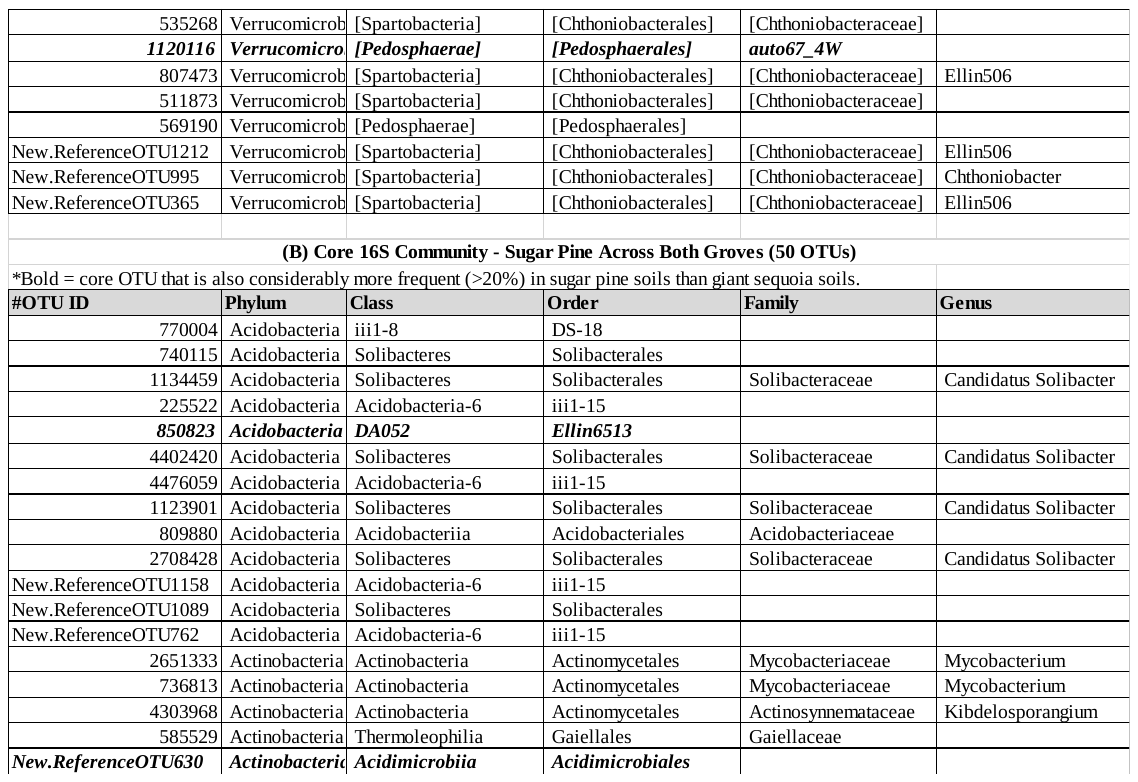

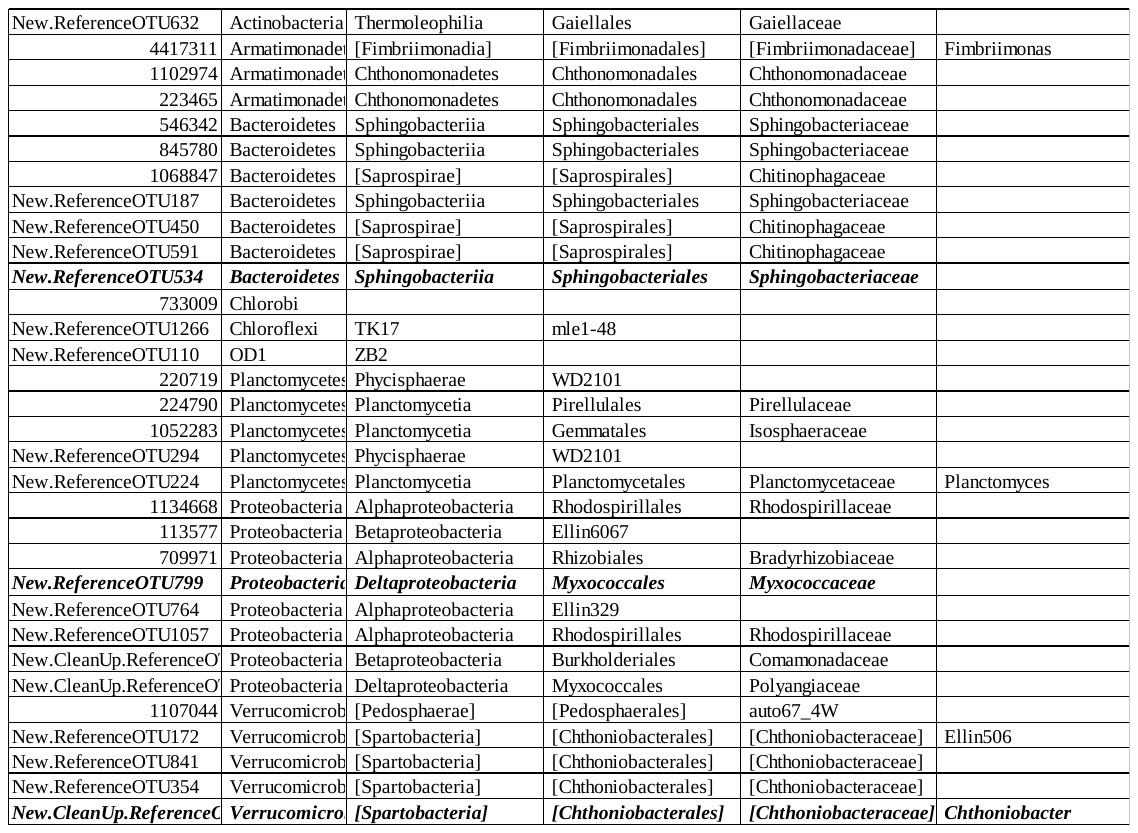

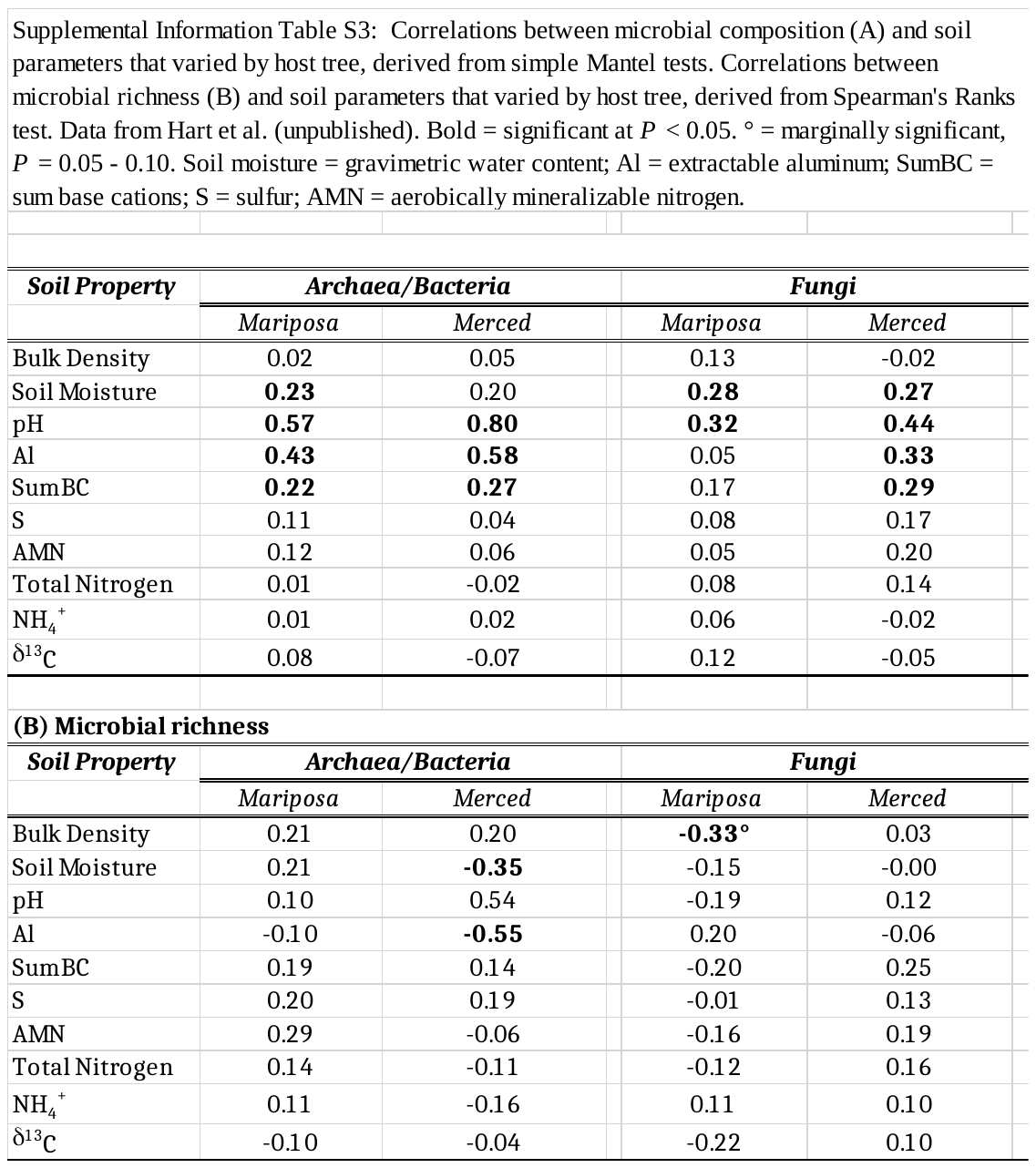
